# DNA demethylases constrain multigenerational methylome and transcriptome divergence

**DOI:** 10.64898/2026.10.29.762914

**Authors:** Anna D. Dmitrieva, Kate I. Christensen, Ben P. Williams

## Abstract

In plants, animals and fungi, DNA methylation is important for the silencing of repetitive DNA and impacts the expression of a subset of genes. DNA methylation landscapes exhibit a high degree of stability over generations, yet this stability is the product of multiple dynamic methylation and demethylation pathways. How these dynamic pathways function over longer multigenerational timescales is unclear. To address this, we inbred quadruple DNA demethylase Arabidopsis mutants and closely related wild type lines for nine generations by single-seed descent. In inbred mutant lines, methylation gains accumulated across generations, with regions of preexisting methylation becoming progressively reinforced or spreading to proximal sequences. Additionally, each inbred line possessed hundreds of de novo methylation gains in unmethylated regions. Two-thirds of all methylation gains and 85% of de novo methylated loci were unique to only one inbred line, suggesting a high degree of stochasticity in multigenerational methylation dynamics. Individual inbred lines also exhibited substantial divergence in both their transcriptomes and small RNA profiles. Our data lead to a model in which DNA demethylases constrain the epigenome and transcriptome space accessible over multigenerational timescales. Long-term loss of DNA demethylase activity causes lineages to drift further apart to more diverged epigenetic landscapes.

**Author summary:** DNA methylation is a chemical mark added to DNA that can impact gene expression. The patterns that DNA methylation marks form across DNA are maintained by enzymes that add and remove them. We wanted to understand the role that DNA methylation removal plays in keeping methylation patterns stable over generations. We followed several genetically identical lineages of Arabidopsis, a small flowering plant, over nine generations. In plants lacking four enzymes that remove DNA methylation marks, methylation gradually accumulated over time. Some existing patches of methylation spread into neighboring regions, and new patches appeared in places that normally lack DNA methylation altogether. Different lineages accumulated mostly different methylation patterns, suggesting that where DNA methylation appears over time is mostly down to chance. Independent lineages also differed in their gene expression patterns, with thousands of genes showing altered expression levels. Our findings suggest that methylation removal limits the range of methylation patterns that can build up over generations in plants. Preserving inherited methylation patterns therefore requires a mechanism of methylation removal, which prevents new marks from taking hold in the wrong places.

## Introduction

Across diverse eukaryotes, DNA methylation provides a critical layer of epigenetic regulation. DNA polymerase cannot incorporate methylated bases into newly synthesized strands. DNA methylation patterns therefore require continual maintenance by DNA methyltransferases [1]. Despite this, DNA methylation landscapes can be stably propagated over 100 to thousands of years [2]. Mutation accumulation lines in *Arabidopsis thaliana* (Arabidopsis), maintained for 30 generations, showed that over 99.998% of the methylated regions remained stably inherited between generations [3]. Such stability in the genomic distribution and levels of DNA methylation enables it to accurately perform its key functions: silencing transposons, regulating gene expression, and establishing imprinting [4]. However, this long-term stability does not reflect biochemical inactivity or a lack of arising epimutations but rather results from continual active maintenance of stable epigenetic states by specialized pathways, a concept called “epigenetic homeostasis” [1].

In plants, DNA methylation occurs at cytosines in CG, CHG, and CHH sequence contexts (H = A, T, C). Methylation is established de novo in all three contexts by the RNA-directed DNA methylation (RdDM) pathway, in which methylation is guided to specific genomic targets by small RNAs (sRNAs) [5]. Once established, methylation is maintained primarily by context-specific methyltransferases, while CHH methylation must be continually re-established *de novo* either by CMT2 (primarily active in heterochromatin) or RdDM (primarily active in euchromatin) [4,6]. Methylation maintenance is aided by self-reinforcing interactions between DNA methylation, the H3K9me2 histone modification, and small interfering RNAs (siRNAs) promoting repeated recruitment of methylation machinery to the same loci [7]. Although these feedback mechanisms are essential for continued silencing of transposons and other repetitive sequences, they can also promote ectopic methylation or methylation spreading from the established targets into neighboring DNA, including genes and their regulatory regions [8]. Mechanisms that remove excessive methylation are therefore essential for enforcing methylation boundaries and protecting gene regulation.

Active DNA demethylation contributes to the maintenance of epigenetic homeostasis by counteracting the activities of DNA methyltransferases. In Arabidopsis, it is performed by four DME-family 5-methylcytosine DNA glycosylases: DEMETER (DME) [9], REPRESSOR OF SILENCING 1 (ROS1) [10], and DEMETER-LIKE 2 and 3 (DML2/3) [11], which prevent genome-wide hypermethylation. The *ros1* mutant and *rdd* (*ros1 dml2 dml3*) triple mutant show thousands of hypermethylated loci, with many demethylase targets also targeted by RdDM [11–13]. To assess the contribution of DME to maintaining genomic methylation patterns in somatic tissues, we previously generated a somatic quadruple *drdd* mutant by rescuing *DME* expression specifically in the central cell, where its expression is required for reproductive development [14]. Methylome analysis in the *drdd* quadruple mutant showed exacerbated hypermethylation compared to the *rdd* triple mutant, including previously undetected hypermethylation inside gene bodies [14].

The trajectory, predictability, and effects of methylation change during prolonged loss of active demethylation remain underexplored. Further, it is unknown whether hypermethylation arises predominantly through reinforcement and expansion of previously methylated regions or through *de novo* methylation gains. A previously published multigenerational inbreeding experiment profiled the *ros1* methylome in generations 1 and 10 and found an increased rate of epimutation accumulation compared to wild-type (*DRDD*, hereafter referred to as “WT”) lines [15]. In this study, we assess the role of DNA demethylases in the long-term stability of DNA methylation inheritance by propagating multiple *drdd* and WT lineages for 9 generations by single-seed descent. We performed methylation profiling at multiple generations, as well as transcriptome and small RNA (sRNA) profiling at the final (9th) generation. Our results show that independent lines accumulate lineage-specific gains of ectopic methylation, leading to divergent methylomes and transcriptomes. New hypermethylation is almost entirely driven by RdDM-pathway activity, occurring both proximally to existing RdDM targets and de novo in unmethylated regions. Our study therefore uncovers long-term RdDM dynamics and the epigenomic consequences of DNA demethylation over multiple generations of inheritance.

## Results

### Inbred quadruple demethylase mutants gain methylation at thousands of intergenic sequences

In this study, three *drdd* and three WT segregant lines [14] were propagated by single-seed descent up to generation 9 (Fig. 1A, see Methods). We hypothesized that, without the homeostatic activity of DNA demethylases, the *drdd* lines would progressively gain methylation over generations. To test this, we performed enzymatic methyl-seq (EM-seq) at the 5th, 7th and 9th inbred generations and identified differentially methylated regions (DMRs) relative to a generation 2 ancestor (see Methods). One of the *drdd* lines had to be terminated after generation 7 as it gradually developed increasingly detrimental phenotypes, with few to no seeds produced. Inbred *drdd* lines showed a clear progressive increase in CG and CHG hypermethylated regions (hyperMRs) over time, with the number of hyperMRs vs the 2nd generation increasing in later generations (Fig. 1B). Conversely, WT inbred lines gained on average ∼6.6 times fewer CG and ∼16.6 times fewer CHG hyperMRs than *drdd* by the 9th generation. Inbred *drdd* lines also had some hypomethylated regions (hypoMRs). In the CG and CHG contexts, there were 15.5-fold and 9.5-fold fewer hypoMRs than hyperMRs, respectively, in the 9th generation (Fig. S1).

**Figure 1.**
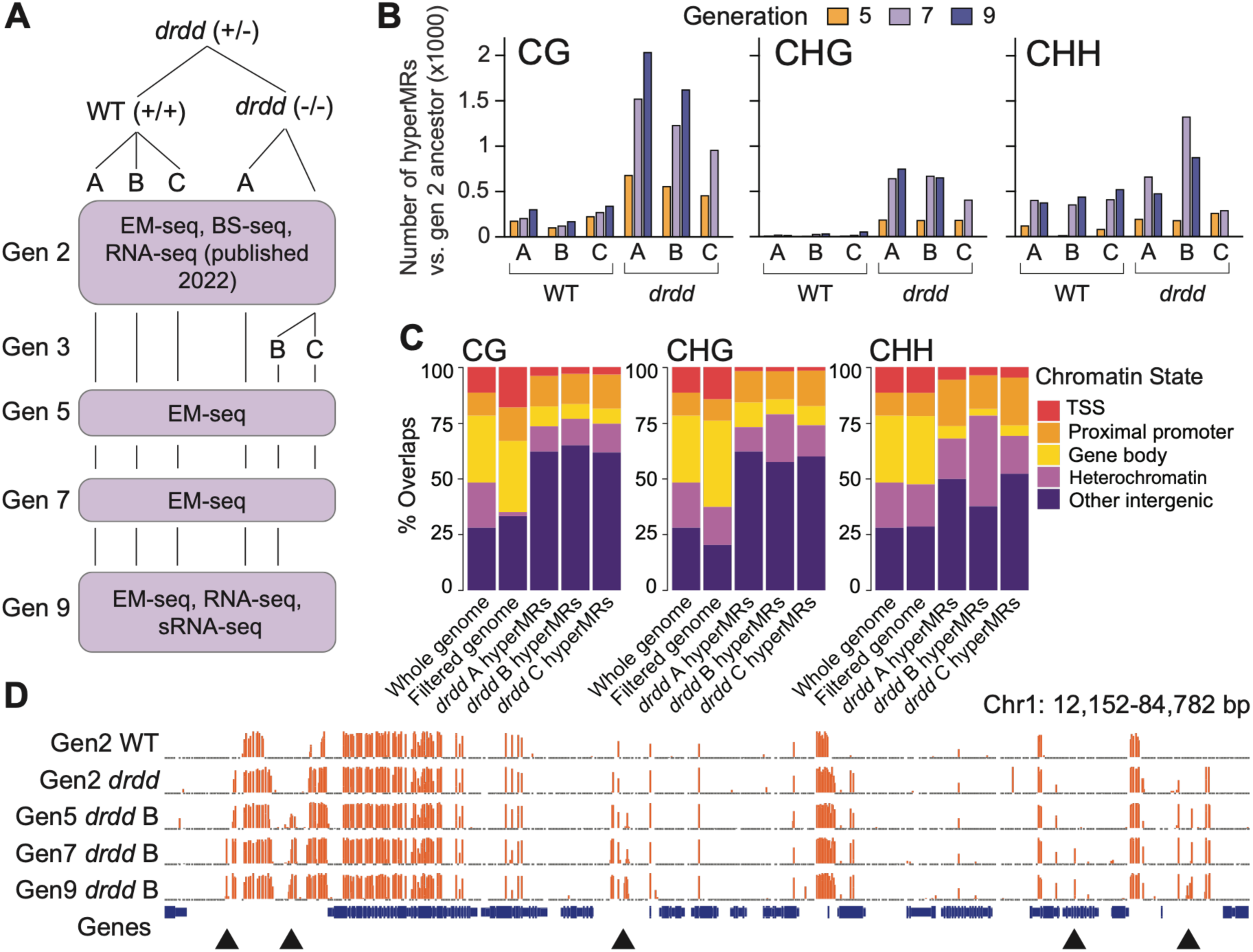
Inbred quadruple demethylase mutants gain methylation at thousands of intergenic sequences. **A.** Outline of the inbred lineages and the sequencing performed. All sequencing was done on whole-leaf tissue. **B.** The numbers of hyperMRs in CG, CHG, and CHH sequence contexts in each line relative to their respective generation 2 ancestor, across all analyzed generations. **C.** Proportions of *drdd* hyperMRs in generation 7 overlapping major chromatin states (see Methods). “Filtered genome” consists of 300 bp bins which have average methylation below 65% in the CG context, 75% in the CHG context, and 90% in the CHH context in generation 2 *drdd* (see Methods). **D.** A representative genome browser snapshot showing regions of progressive methylation gain in the CG context between a generation 2 *drdd* ancestor and three later generations of *drdd* line B. A generation 2 WT sample is shown as a control. Black arrowheads indicate sites of methylation gain relative to generation 2 *drdd*. The vertical scale for each track ranges from 0 to 100% methylation. Thin dark gray bars beneath each track indicate CG sites with EM-seq coverage but no detected methylation (0%).

We next wanted to assess the chromatin landscapes where hyperMRs are most prevalent. We condensed previously characterized chromatin states [16] into five categories and assessed the proportion of hyperMRs overlapping each. In WT inbred lines, hyperMRs were most enriched in gene bodies for CG methylation, consistent with the known patterns of epimutation described in WT inbreeding experiments [17] (Fig. S2). Additionally, CHG and CHH hyperMRs in WT were most enriched in heterochromatin (Fig. S2). In contrast, hyperMRs in all three *drdd* inbred lines were more strongly enriched in non-heterochromatin intergenic regions and promoter regions (Fig. 1C-D). This is substantially different from both the distribution of naturally occurring epimutations that occur in WT, as well as the genomic distribution of chromatin states. Together, these analyses demonstrate that inbreeding *drdd* results in the accumulation of a large number of methylation gains that are mostly distinct from the natural baseline processes of epimutation.

Because several DNA methylation pathways rely on self-reinforcing loops, we wanted to assess whether the methylation gained in *drdd* over the course of inbreeding was largely spreading from existing regions of methylation. We assigned hyperMRs to four classes based on their location relative to methylated regions in a second-generation ancestor methylome (see Methods). These classes are: “contained” for hyperMRs fully overlapping previously methylated regions, “spreading” for hyperMRs located <= 600 bp immediately adjacent to ancestral methylation, “de novo*”* for hyperMRs >1200 bp away from previous methylation in any context, and “ambiguous” for hyperMRs not matching any of the previous conditions (Fig. 2, A-B). Between 50.8 and 74.3% of all *drdd* hyperMRs in generation 7 were classified as either “contained” or “spreading,” suggesting methylation gains at existing methylated regions (e.g. a higher percentage of methylated reads compared to the ancestor), or gains at cytosines immediately adjacent to ancestrally methylated regions (Fig. 2C). However, we also identified 14.1–31.2% of all generation 7 *drdd* hyperMRs as de novo (Fig. 2C), occurring in regions lacking DNA methylation in the ancestor. Notably, de novo hyperMRs were very rare in inbred WT control lines, suggesting their presence is dependent on the loss of demethylation by DRDD. These de novo hyperMRs in *drdd* are completely absent in heterochromatin and have a higher enrichment in non-heterochromatin intergenic regions compared to the genomic distribution of all hyperMRs (Fig. S3).

**Figure 2.**
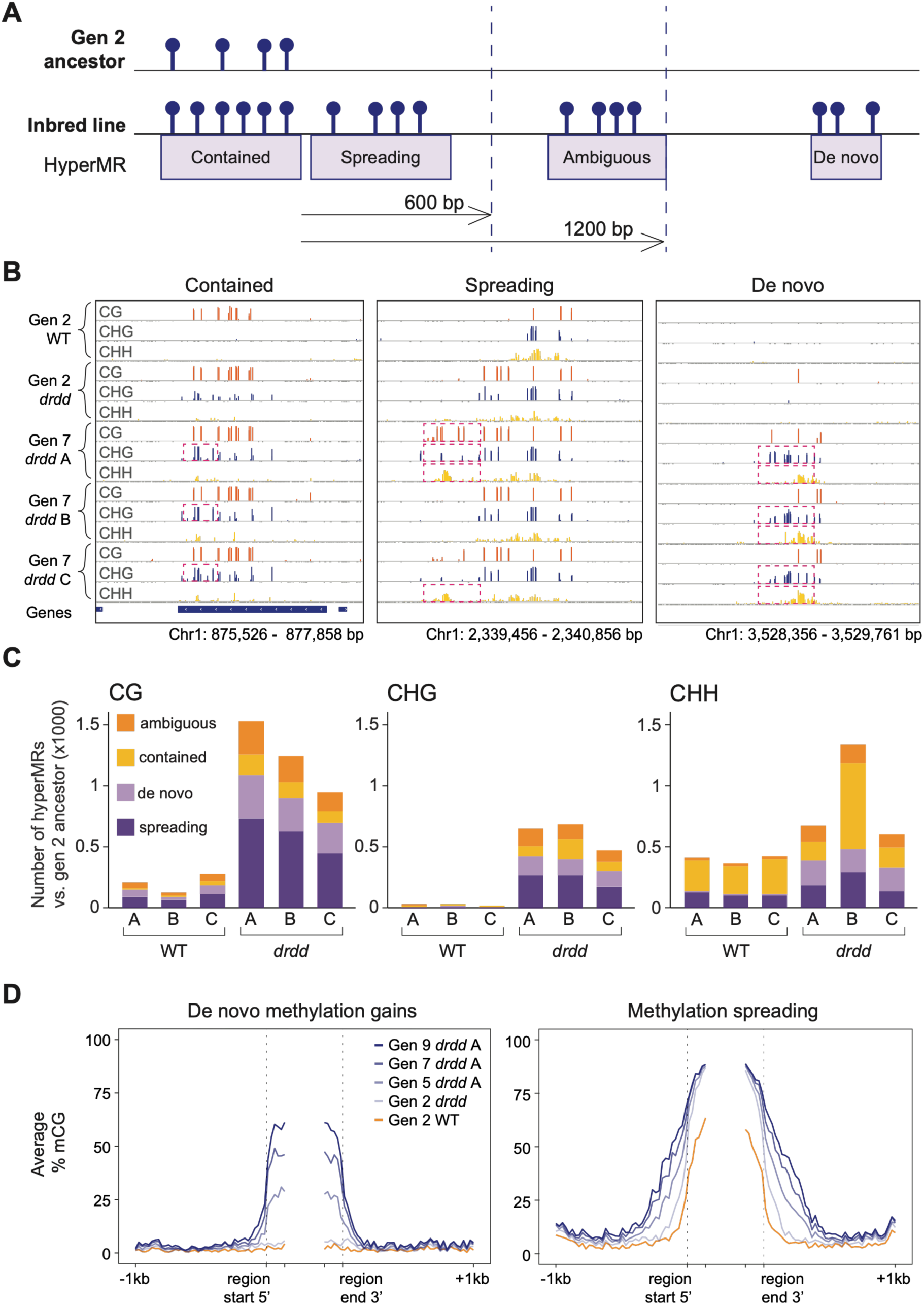
De novo and spreading methylation gains happen progressively over the course of inbreeding. **A.** The distance thresholds used for classifying hyperMRs. Distances are measured in base pairs from methylated regions which have conserved methylation between the generation 2 ancestor and an inbred line. **B.** Representative “contained”, “spreading”, and “de novo” methylation gains in *drdd* lines in generation 7, in all sequence contexts. For the “contained” class, mCHG is elevated in generation 7 relative to generation 2 *drdd*. Generation 2 WT is shown as a control. Dashed pink outlines show called hyperMRs. The vertical scale for each track ranges from 0 to 100% methylation. Thin dark gray bars beneath each track indicate sites with EM-seq coverage but no detected methylation (0%). **C.** Numbers of hyperMRs in generation 7, showing their distribution across the four classes. **D.** Ends analysis plots showing average CG methylation levels over de novo and spreading hyperMRs called in generation 9 in *drdd* line A, relative to generation 2 *drdd.* Generation 2 WT is shown as a control. 150 bp are shown on either side inside the dashed lines, and methylation is averaged over 25 bp bins. Left: de novo hyperMRs are between the vertical dashed lines. Right: spreading hyperMRs are outside of the vertical dashed lines, and the ancestral methylated regions adjacent to them are between the dashed lines.

To understand the stability and fate of acquired hyperMRs, we assessed methylation profile dynamics at the hyperMRs over generational time. We examined the methylation profiles of *drdd* lines at each generation over generation 9 hyperMRs to trace how their methylation changed over time (Fig. 2D). Both de novo and spreading hyperMRs exhibited progressively increasing methylation levels from generation 2 onwards through inbreeding (Fig. 2D, S4). Consistently, de novo hyperMRs exhibited an increased average percentage of methylation over generations, whereas spreading hyperMRs exhibited a clear increase in the spreading distance as generations progressed (Fig. 2D). This suggests that once gained, these newly methylated regions were inherited and/or reinforced in subsequent generations.

### De novo methylation gains are RNA-directed

Methylation spreading outwards from previously methylated regions in DNA demethylation mutants is a previously described phenomenon attributed to RdDM activity [11,18,13]. To examine if RdDM could also be the mechanism underpinning the formation of spontaneous de novo hyperMRs, we assessed the sequence contexts of methylation gains in these regions. We found that on average 73.7% of all *drdd* de novo hyperMRs in generation 7 had a methylation gain of 5% or more in at least one other context (Fig. 3A). As multi-context methylation gains are a hallmark of RdDM activity, we hypothesized that sRNAs are likely being produced to target methylation to these regions. To test this, we performed sRNA-seq on the 9th inbred generation of *drdd* and WT control lines. We found that most frequently, *drdd* de novo hyperMRs were associated with *drdd*-specific sRNA clusters which are absent in WT lines, while other hyperMR classes had much higher overlap with WT sRNA clusters as well (Fig. 3, B-C). This relationship was especially strong for CHH *drdd* de novo hyperMRs where 81.4-91% were associated with new sRNA clusters. Canonical RdDM relies primarily on 24-nucleotide (nt) siRNAs as guides and also generates 23 nt siRNAs [19,20]. Indeed, on average 92.8% of the sRNA reads underlying *drdd* de novo hyperMRs are 23–24 nt long, with around 74.7% being 24 nt (Fig. 3D). These data therefore strongly suggest that the RdDM pathway is the primary mechanism contributing to de novo methylation gains in inbred *drdd*.

**Figure 3.**
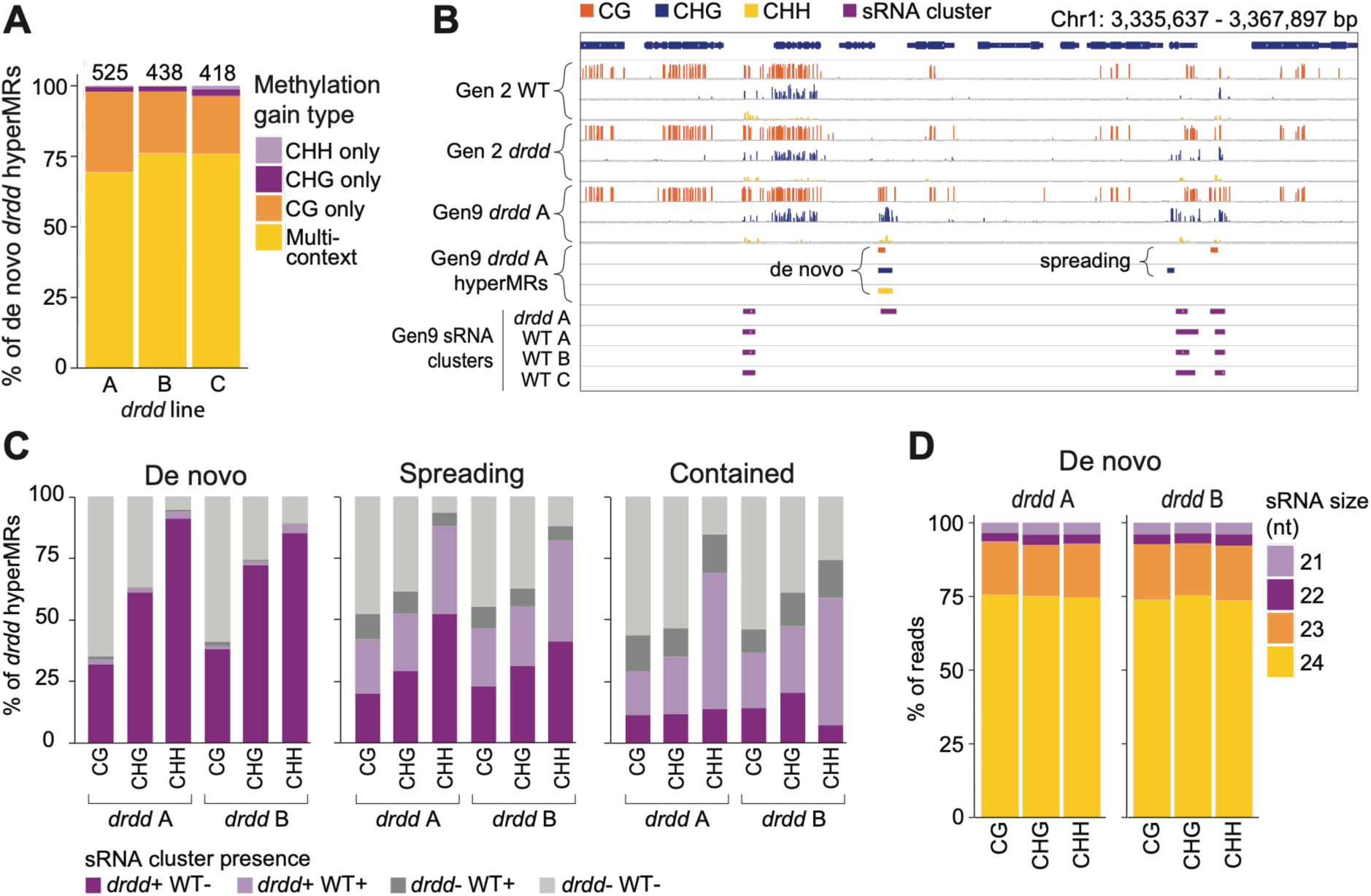
De novo methylation gains are RNA-directed. **A.** Distribution of multi-context and single-context methylation gains in generation 7 *drdd* lines at supersets of de novo hyperMRs in all contexts. Multi-context gains were defined as de novo hyperMRs that also exhibited an average methylation gain of at least 5% in at least one additional sequence context. **B.** A representative genome browser snapshot showing de novo and spreading hyperMRs in generation 9 *drdd* line A and underlying sRNA clusters. The vertical scale for each methylation track ranges from 0 to 100% methylation. Thin dark gray bars beneath each track indicate sites with EM-seq coverage but no detected methylation (0%). **C.** Distribution of generation 9 *drdd* hyperMRs by overlap with sRNA clusters in the corresponding *drdd* line and in any WT line. **D.** Distribution of sRNA read lengths for sRNAs underlying de novo hyperMRs in 9th generation *drdd* lines.

### De novo hyperMRs occur in regions associated with Pol V activity in wild type Col-0

In addition to siRNAs, RdDM relies on long noncoding RNA scaffolds that enable recruitment of complementary AGO4-bound siRNAs to loci targeted for DNA methylation. These RNA scaffolds are synthesized by a specialized plant RNA polymerase, polymerase V (Pol V) [5]. Recently, low levels of Pol V activity have been detected at a large portion (potentially as high as 30%) of the wild type Col-0 genome [21], including many “non-RdDM” regions that are not associated with sRNAs or non-CG methylation. We therefore reasoned that these regions of the genome exhibiting Pol V activity may have a higher probability of nucleating new RdDM sites in inbred *drdd*. We intersected each hyperMR class with previously published regions predicted to have Pol V transcription in Col-0 [21]. More than two thirds of de novo hyperMRs overlapped non-RdDM Pol V-transcribed regions (Fig. 4A-B), supporting the hypothesis that pre-existing wild type Pol V activity in these regions may prime them for de novo methylation in *drdd* mutants. Removal of methylation by DRDD may explain the lack of RdDM occurring at these sites in wild type Col-0. As expected, contained hyperMRs (which directly overlap previously methylated regions) and spreading hyperMRs (which are adjacent to methylated regions) were associated with regions of Pol V activity that are identified as RdDM targets in Col-0 (Fig. 4A). These data suggest that Pol V activity may be one of the key factors that predisposes a region to gain of new RdDM activity in *drdd* mutants, and that regions with Pol V activity already detectable in wild type Col-0 are likely locations for hypermethylated regions during *drdd* inbreeding.

**Figure 4.**
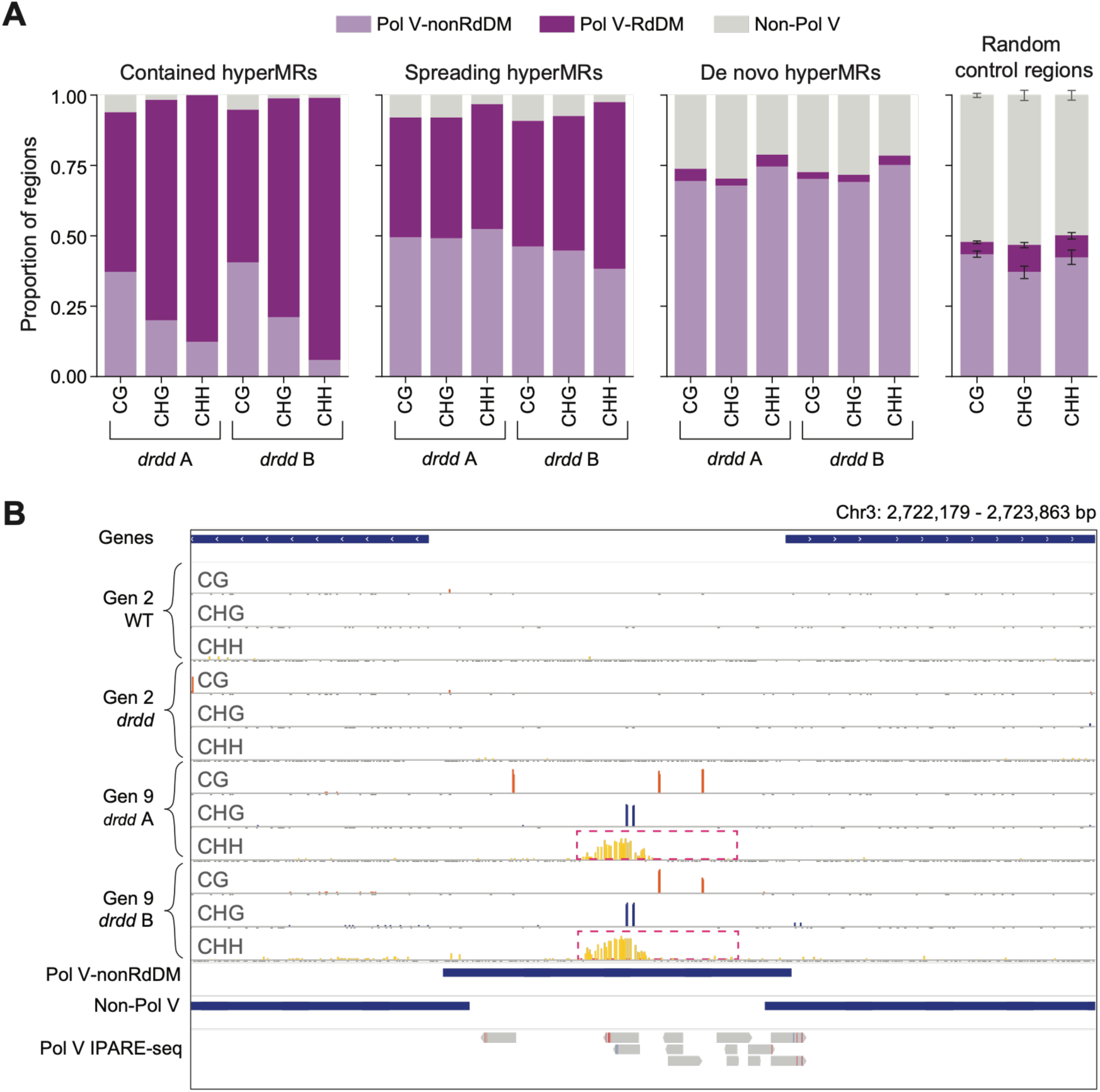
De novo hyperMRs occur in regions associated with Pol V activity in wild type Col-0. **A.** Distribution of hyperMRs of each class in generation 9 by overlap with previously published predicted wild type Col-0 Pol V transcription states [21]. “non-Pol V” is a combination of regions predicted to be untranscribed and transcribed by other polymerases. Random controls are 300 bp regions which were lowly methylated in generation 2 *drdd* (average mCG < 50%, mCHG < 20%, and mCHH < 5%). Error bars indicate standard deviation, calculated from five independent random subsamples of the random control regions. **B.** A representative genome browser snapshot showing a CHH de novo hyperMR called in both *drdd* lines in generation 9, with underlying Pol V IPARE-seq reads in wild type Col-0. Predicted WT Col-0 Pol V transcription states in this region are shown as horizontal dark blue bars, and dashed pink outlines show called de novo hyperMRs.

### Loss of DRDD leads to long-term methylome and transcriptome divergence

Given that multiple independent *drdd* lines had a similar distribution of hyperMR classes (Fig. 2C), similar underlying RNA distributions (Fig. 3C) and Pol V activity in WT (Fig. 4A), we predicted that independent *drdd* lines may share many common hyperMRs and exhibit predictable gains of methylation at the same loci. To test this hypothesis, we plotted average methylation over a superset of all hyperMRs ever called, between all lines and generations. In contrast to our initial hypothesis, methylation levels at all hyperMRs across different *drdd* lines showed distinct patterns (Fig. 5A, S5), suggesting that methylation was initiated at unique sites between the lines and further amplified to form progressively divergent methylomes. Consistently, 65.7–85.4% of all hyperMRs were unique to one *drdd* line, with de novo hyperMRs being the most line-specific (85.0–85.4% unique) (Fig. 5B, S5). Additionally, most *drdd*-specific sRNA clusters underlying hyperMRs were unique to one of the two *drdd* lines profiled (Fig. S7). In contrast, WT lines retained stable average methylation at the *drdd* hyperMRs through the course of inbreeding (Fig. S6).

**Figure 5.**
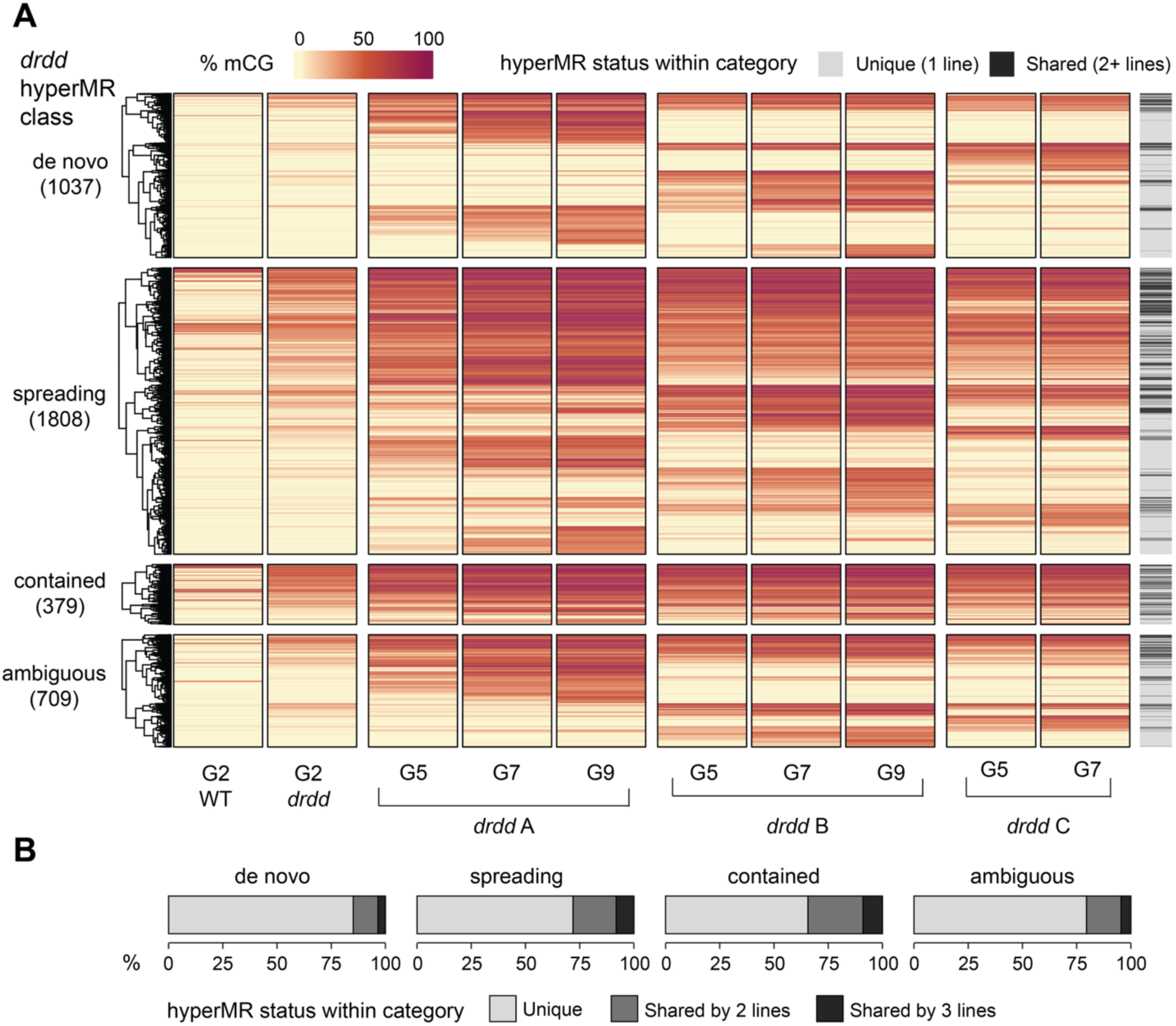
*drdd* lines have divergent methylomes. **A.** Heatmap of average methylation in *drdd* lines over class-specific supersets of all CG hyperMRs ever called, between all lines and generations. Generation 2 WT is shown as a control. HyperMRs are sorted by hierarchical clustering within each class. Total numbers of CG hyperMRs in each reduced superset are included in parentheses underneath each hyperMR class name. In the hyperMR status within each class, “Unique (1 line)” means a hyperMR was only called in one line throughout the experiment. “Shared (2+ lines)” means a hyperMR ever called in one line was called in at least one other line in at least one of the generations. **B.** Distribution of *drdd* hyperMRs present in 1 line or shared between 2 or 3 lines in each hyperMR class.

Because hyperMRs in the inbred *drdd* were enriched in non-heterochromatic intergenic regions and promoters (Fig. 1C), we hypothesized that they may be significantly affecting gene expression. We performed RNA-seq on inbred *drdd* and WT lines in the 9th generation, to assess the effects of the highest final numbers of accumulated hyperMRs in *drdd* on transcription, as well as the impacts of multigenerational methylome divergence. We performed RNA-seq using six biological replicates of each WT and *drdd* line and called differentially expressed genes (DEGs) separately for each of the two *drdd* lines. First, we compared the numbers of DEGs to the generation 2 *drdd* [14] and generation 4 *drdd* [22]. The *drdd* lines in generation 9 showed a substantial increase in the numbers of DEGs, with a total of 4,057 and 2,539 DEGs in *drdd* lines A and B, respectively, ∼6-9.6 times higher than in generation 2 and ∼3.3-5.3 times higher than in generation 4 (Fig. 6A). Identifying DEGs with a reduced randomly subsampled number of WT replicates did not impact this result (S3 Table).

**Figure 6.**
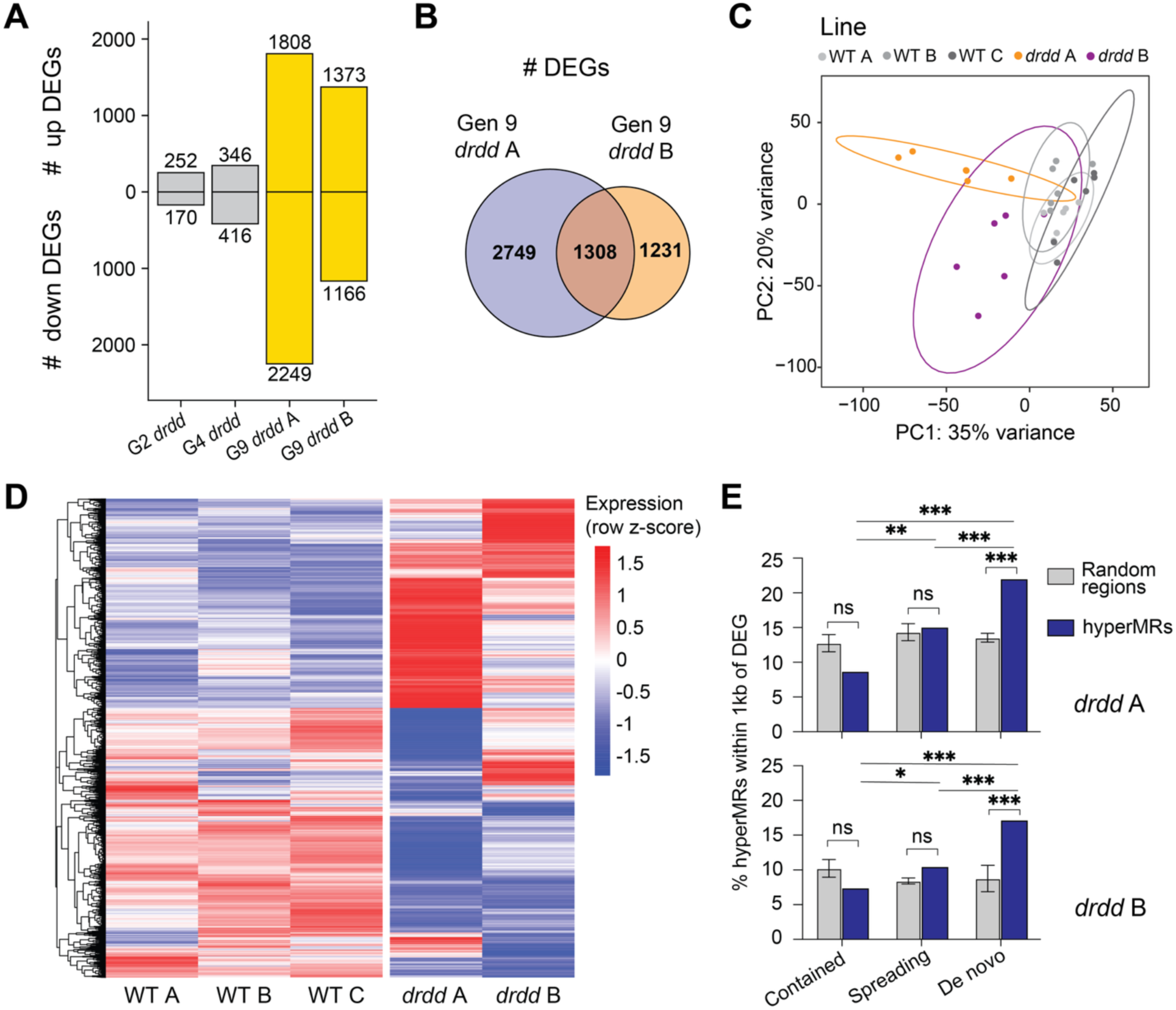
*drdd* lines have divergent transcriptomes most strongly associated with de novo hyperMRs. **A.** Numbers of DEGs in the two generation 9 *drdd* lines relative to WT lines, compared to those previously reported for generations 2 and 4 in separate studies [14,22]**. B.** Overlap between genes separately classified as DEGs relative to WT lines in the two generation 9 *drdd* lines. **C.** PCA of rlog-transformed gene-expression counts for all genes, with ellipses indicating 95% multivariate *t*-distribution data regions. **D.** Heatmap of row-scaled normalized expression across the union of *drdd* DEGs relative to WT. **E.** Percentages of hyperMRs of each class in generation 9 located within 1 kb of a DEG. Random controls consisted of randomly selected 300-bp regions matched in number to the hyperMRs in each corresponding class. Error bars indicate standard deviation, calculated from five independent random subsamples of the random control regions. Significance between hyperMR classes was tested using the Chi-squared test, and significance between random windows and hyperMRs was tested using Fisher’s exact test. For P-value cutoffs, * indicates P < 0.05, ** indicates P < 0.01, and *** indicates P < 0.001.

The two independently profiled 9th generation *drdd* lines shared 1,308 DEGs, suggesting they do not share a closely related set of transcriptome changes (Fig. 6B). Principal component analysis (PCA) showed that the two *drdd* lines occupied largely distinct regions of transcriptional space from one another, as well as from the WT lines, which clustered more closely together (Fig. 6C). Consistent with this pattern, a heatmap of normalized expression across the union of all *drdd* DEGs showed broadly similar expression profiles among the WT lines but distinct expression patterns in the two *drdd* lines (Fig. 6D). These data suggest that the divergence in methylation patterns between the two lines is associated with divergent transcriptome changes. Consistently, direct comparison of the two *drdd* lines identified 2,641 DEGs between them, further supporting their substantial transcriptional divergence. To examine the biological processes associated with these divergent transcriptional profiles, we performed Gene Ontology (GO) enrichment analysis separately for the DEGs identified in each *drdd* line. Both lines showed enrichment for defense- and specialized metabolism-related processes among the top enriched terms. Additionally, line A had line-specific enrichment for translation, ribosome biogenesis, and cell-cycle-related processes (Fig. S8).

To examine if the methylation gains caused by multigenerational loss of DRDD activity may underpin some of these transcriptome changes, we examined the proximity between hyperMRs and DEGs called in each independent *drdd* line. We observed that a small but nevertheless significant subset of hyperMRs are in close genomic proximity (≤ 1 kb) to DEGs (Fig. 6E), similar to the association between DMRs and DEGs previously described [14]. Furthermore, this analysis showed that out of all hyperMR classes, de novo hyperMRs are more frequently associated with DEGs, and therefore may be the primary driver of transcriptome changes, with 17–22% of de novo hyperMRs being located within 1 kb of a DEG (Fig. 6E). Together, this multigenerational multi-omic profiling of inbred lines leads to a model in which the loss of DRDD leads to a shift in the accessible epigenetic landscape over time. In WT plants, DNA demethylation is restricting the extent to which newly methylated regions can form and consequently constraining the extent to which methylomes can diverge from one another within related individuals. When the DRDD pathway is knocked out, independent lines are able to drift further apart across epigenetic space, leading to divergent methylation patterns and substantially different gene expression profiles.

## Discussion

In this study, we demonstrate that the DNA demethylases in Arabidopsis stabilize the methylome over multiple generations of inheritance by suppressing de novo methylation across the genome. Our study also shows that in the absence of DNA demethylase activity, de novo methylation events are stochastic and specific to individual lineages, causing independent lineages to diverge in epigenetic space along distinct trajectories. While hypermethylated loci are distinct between individual *drdd* mutant lines, they appear to be drawn from a constrained subset of the genome that is susceptible to methylation gain. The creation of distinct epigenomic states in individual inbred lines is accompanied by substantial transcriptional divergence, suggesting that the methylation gains prevented by DNA demethylation activity can broadly impact gene regulation programs. Stable epigenetic inheritance therefore depends not only on faithful propagation of existing methylation patterns, but also on the continual prevention of ectopic methylation from becoming established and amplified by strong positively reinforcing methylation pathways.

Our data strongly suggest that RdDM is responsible for the majority of methylation gains occurring in inbred *drdd* mutants. While we observe the hallmarks of RdDM maintaining methylation at these sites, such as the presence of 24 nt sRNAs, these snapshots do not fully explain how such de novo methylation events are initiated in an unmethylated parent genome. While methylation at RdDM targets is reinforced through the canonical Pol IV-dependent pathway, de novo methylation can be initiated through several noncanonical RdDM pathways in which 21–22 nt sRNAs are generated from a variety of other sources, including aberrant transcripts [23,24]. Notably, most of the de novo sRNA-targeted hyperMRs we observed were in typically non-transcribed (by RNA polymerase II) intergenic regions. HyperMRs are therefore either initiated from non-polymerase II sources, such as RNA polymerase IV, or are the product of low-frequency polymerase II activity outside of typically transcribed regions.

Our data suggest that some genomic regions are more susceptible to de novo methylation than others. We observe de novo methylation events more frequently in certain chromatin states (Fig. S3), and a subset of loci became methylated in multiple independent lines (Fig. 5). A central question therefore is – why are some loci more susceptible to de novo methylation gain than others? The lineage specificity of many *drdd* de novo hyperMRs and their frequent overlap with regions transcribed by Pol V in WT [21] support a “partial randomness” model in which de novo methylation is established stochastically within a constrained subset of susceptible loci. This association is consistent with the proposed genome-surveillance model in which low-level Pol V transcription makes genomic regions competent for RdDM if complementary siRNAs become available [21]. Some evidence suggests that siRNA-guided ARGONAUTE recruitment precedes and directs Pol V recruitment, in which case Pol V activity may not reflect a genomic surveillance function but rather a signature of low-frequency RdDM activity at certain genomic regions [25]. Once initial methylation became established at the new RdDM targets in *drdd*, SUVH2 and SUVH9 could facilitate continued Pol V recruitment, creating a positive feedback mechanism and contributing to RdDM reinforcement and the formation of newly methylated regions [26].

In addition to the gain of many newly methylated regions, inbreeding also causes substantial transcriptomic divergence over time. Inbred *drdd* mutant lines exhibit thousands of DEGs, substantially more than the number identified in non-inbred *drdd* mutants. In both mammals and plants, only a subset of genes are sensitive to proximal DNA methylation changes [27,28] and genome-wide correlations between methylation and expression changes are typically weak. Our data are consistent with this, with a minority of DEGs correlating with proximal hyperMRs that occurred during inbreeding, typically within promoter regions. While our data cannot distinguish between direct effects of proximal methylation changes and indirect effects from broader transcriptional networks, the fact that transcriptomes were substantially different in independent lines with divergent methylomes suggests that the transcriptomic changes are a consequence of methylation changes in gene regulatory regions. Notably, at the organismal level, several *drdd* lineages became non-viable and could not be maintained through the full experiment, suggesting that different methylation trajectories potentially produce different fitness consequences. The lineages analyzed at generation 9 therefore represent a higher-fitness subset of the possible outcomes of prolonged demethylation loss, potentially causing us to underestimate its most severe epigenomic and transcriptional consequences.

Our study demonstrates that the activity of DNA demethylases is essential for stable epigenetic inheritance over multiple generations. This mirrors the finding that elevated expression of a DNA demethylase also causes unstable inheritance of methylation over generations [29]. DRDD activity must therefore be neither too high nor too low in order to enable optimal propagation of epigenetic states and constrain epigenome divergence over generations. Mechanisms such as the DNA methylation rheostat (methylstat) at the promoter of *ROS1* [30,31] are key to balancing demethylation activity with the need for long-term stable epigenetic inheritance.

## Materials and Methods

### Plant inbreeding and growth conditions

The seeds from second generation individual *drdd* and WT segregant progenitor plants were obtained from Williams et al. [14]. Two of the four *drdd* lines already maintained to the third generation showed detrimental phenotypes, with plants that became sterile within 1–2 additional generations, and these lines were excluded from the study. To maintain three *drdd* lines, we derived two sub-lines from siblings of one parent line at generation 3 (Fig. 1A). The resulting 6 lines (*DRDD* lines A, B, and C; and *drdd* lines A, B, and C) were subsequently inbred through single-seed descent through 9 generations total, with the exception of *drdd* line C which became sterile and had to be terminated after generation 7. At each generation, plants that were selected for genomic profiling were genotyped to confirm the correct genotype for each sample (primers provided in S1 Table). Seeds were stratified for ∼2 days at 4 °C and grown under long-day conditions (LD, 16 h light / 8 h dark, ∼21 °C, ∼60% humidity) in a Percival growth chamber (AR100L3). Most sample collection happened at 21 days after germination (DAG), described in more detail in subsequent sections.

### Enzymatic methyl sequencing

Whole-genome single-cytosine-resolution DNA methylation was profiled with enzymatic methyl sequencing (EM-seq) [32]. For each line at each generation assessed, a single 5th true leaf was sequenced. Whole-leaf tissues were collected, flash-frozen in liquid nitrogen, and stored at −80 °C until use. In the 5th generation, 5th true leaves were collected when the plants were visually at a similar developmental stage between 21–28 DAG. Frozen leaf tissue was processed using a Qiagen TissueLyser II and incubated in CTAB buffer at 65 °C for about 30 minutes, followed by the addition of 3 μl of RNase and incubation at 37 °C for 20 minutes. DNA was then purified with chloroform, precipitated in 100% ethanol at −20 °C, washed with 70% ethanol, and eluted with nuclease-free water. DNA was sheared to an average fragment size of ∼300 bp with a Covaris S220 Focused Ultrasonicator, using 99-second sonication. EM-seq libraries were constructed according to the NEBNext® Enzymatic Methylseq Kit protocol with either 25 or 50 ng starting DNA, depending on the sample DNA concentration. Library quality was assessed using a KAPA Library Quantification Kit, and average fragment size was verified using a fragment analyzer. Libraries were sequenced with Illumina NovaSeq X, with 150 bp paired-end reads.

EM-seq data were mapped as described in Williams et al. [14]. Briefly, adapters were removed using Trim Galore 0.6.6 (Babraham Bioinformatics). Filtered reads were mapped to the *Arabidopsis thaliana* TAIR10 genome using Bismark 0.22.3 (Babraham Bioinformatics), allowing for 0 mismatched reads in the seed region and removing PCR duplicates. Percent methylation for each cytosine was calculated using the Bismark methylation extractor function. The EM-seq conversion efficiency was calculated based on the percentage of methylation of reads mapped to the chloroplast, which is unmethylated. Conversion rates were > 98.7% across all samples sequenced. Genome browser snapshots for EM-seq data and all other genomic data were taken in Integrative Genomics Viewer (IGV) v. 2.16.0 [33]. Sequencing and mapping statistics for all EM-seq libraries are provided in S2 Table.

### DMR analysis

Differentially methylated regions (DMRs) were called using DMRcaller [34]. DMRs were called between pairs of samples: a generation 2 ancestor and an inbred line sample. Lists of hyperMRs and hypoMRs are provided in S1–S4 Data. The generation 2 ancestors used are a *drdd* plant and a *DRDD* plant, which are siblings of the generation 2 line progenitor plants profiled in Williams et al. [14]. Before using DMRcaller, read counts were first corrected based on conversion rate as previously described [35]. DMRcaller was run using the bins method, with the following settings: bin size of 300 bp; p-value threshold of 0.01; minimum cytosine count of 10; min proportion difference of 0.35 for CG, 0.25 for CHG, and 0.1 for CHH; min gap between DMRs of 500 bp, and min reads per cytosine of 5. The resulting DMRs were filtered for hyperMRs, as these are most likely to be a direct impact of the mutants losing demethylase activity. To analyze chromatin states, the resulting hyperMRs were overlapped with the nine chromatin states previously defined by Sequeira-Mendes et al. [16]. The nine states were reduced into five chromatin functional groups that were most enriched in the original 9 states: transcription start site (TSS) - state 1; proximal promoter - state 2; gene body - states 3, 6, and 7; heterochromatin - states 8 and 9; other intergenic - states 4 and 5. To generate control chromatin state distributions, the *drdd* and WT generation 2 ancestor genomes were filtered for 300 bp bins that had average methylation below 65% in the CG context, 75% in the CHG context, and 90% in the CHH context to retain only regions which were eligible for being classified as a hyperMR based on DMRcaller cutoffs. These control distributions are shown as “filtered genome” in chromatin state distribution plots, with the *drdd* control distribution used in Fig. 1C and S3 and the WT control distribution in Fig. S2.

To classify hyperMRs based on their location relative to previously methylated regions in generation 2, a custom R script was written. Briefly, the mapped EM-seq sequencing data from generation 2 were broken into 300 bp bins, and average methylation was calculated over each bin. Resulting bins were filtered to retain those with at least 10 cytosines and an average of 5 reads per cytosine. Regions were classified as methylated if the average methylation was at least 50% for CG, 20% for CHG, or 5% for CHH. Methylated bins that were up to two bins (600 bp) apart were merged, unless the merged bin no longer met the cytosine and read requirements, in which case they were kept separate. The output of these steps was a set of “generation 2 methylated” regions. Next, regions that retained methylation were identified in each inbred line at each generation, and regions that either lost methylation, or did not pass the same cytosine count, average reads, and average methylation threshold filters were filtered out, resulting in a final set of “retained generation 2 methylated” regions. Finally, hyperMRs found for each inbred line were classified based on their location relative to retained generation 2 methylated regions: “contained” for hyperMRs fully contained in the original region, “spreading” if the hyperMR overlapped the two closest bins (600 bp upstream or downstream), “de novo” for hyperMRs beyond 1200 bp, and “ambiguous” for any hyperMRs not matching any of the previous conditions, such as falling between the two distance cutoffs. The distances were chosen based on a previous finding that methylation tends to spread up to ∼500 bp away from heavily-methylated siRNA targets [8] and 600 bp being the closest distance that is a multiple of the 300 bp bin length for both DMRcaller and our script.

The heatmap of average methylation across a superset of all *drdd* hyperMRs was generated in R using the package ComplexHeatmap [36], using hierarchical clustering to sort hyperMRs within each class. Within each hyperMR class and methylation context, hyperMRs identified across all *drdd* lineages and generations were combined, and overlapping intervals were merged to generate a nonredundant superset of regions which were used for the heatmap. Intersect analysis for methylation and all other data was performed using bedtools intersect, and proximity analysis was performed using bedtools closest [37]. Out of figures depicting hyperMRs in a single generation or relying on single-generation hyperMRs for the underlying analysis, figures 1C, 2B, 2C, 3A, S2, and S3 are based on hyperMRs in the 7th generation, so as to compare three independent inbred lines. Figures 2D, 3B, 3C, 3D, 4A, 4B, 6E, S4, and S7 are based on hyperMRs in the 9th generation, so as to analyze methylation gains in the final inbred generation or to correlate methylation data with same-generation RNA-seq or sRNA-seq data.

### RNA-seq

The 5th true leaf of six independent replicate plants was collected at 21 DAG for each line analyzed at generation 9. Whole-leaf tissues were collected, flash-frozen in liquid nitrogen, and stored at −80 °C until use. Total RNA was isolated using a QIAGEN RNeasy Plant Mini Kit, following the manufacturer’s protocol. Briefly, frozen leaf tissue was ground to a powder using glass beads using Qiagen TissueLyser II and resuspended in Buffer RLC with β-mercaptoethanol. Plant debris was removed through centrifugation in a lilac spin column, followed by RNA precipitation with 100% ethanol, washes with buffer RW1 and buffer RPE, and elution in RNase-free water performed through centrifugation in a pink spin column.

Library preparation and sequencing were performed by Novogene, with sequencing done on a NovaSeq 6000 using 150-bp paired-end reads. All samples were sequenced in one batch. The reads were mapped as previously described [38]. Briefly, reads were first processed with Trim Galore v0.6.6 (Babraham Bioinformatics) and cutadapt [39]. During this step, adapter sequences were trimmed, the first 9 bp were clipped from the 5’ end of each read, low-quality bases were trimmed from the 3’ end using a Phred quality threshold of 25, and reads shorter than 20 bp after trimming were discarded. Reads were mapped to the Araport11 genome using STAR 2.7.1a [40], and alignments were filtered to retain uniquely mapped reads, allowing mismatches up to 5% of the total read length. Sequencing and mapping statistics for all RNA-seq libraries are provided in S2 Table. Gene-level read counts were generated with htseq-count from HTSeq v2.0 [41] and are provided in S5 Data. Normalization and differential expression analysis were performed using DESeq2 v1.40.1 [42]. Principal component analysis (PCA) was performed on regularized-logarithm (rlog)-transformed gene counts using DESeq2 (S6 Data), and 95% multivariate *t*-distribution ellipses were plotted for each line using ggplot2 [43]. Differential expression was called separately for each of the two *drdd* lines, against a set of all 18 WT replicates (S7 and S8 Data). We also repeated the DEG analysis with 6 WT replicates selected by random subsampling and achieved equivalent numbers of DEGs (S3 Table; S9 and S10 Data). DEGs were defined as genes with log2 fold-change >1 and a Benjamini–Hochberg-adjusted P value < 0.05.

GO enrichment analysis was performed separately for DEGs called in each *drdd* line using the PANTHER 19.0 Statistical Overrepresentation Test [44]. For each analysis, the reference list consisted of all genes with non-NA adjusted P-values in the corresponding DESeq2 comparison (24,003 genes for each line). Enrichment was tested using the GO Biological Process Complete annotation dataset, Fisher’s exact test, and false discovery rate (FDR) correction for multiple testing. Overrepresented terms with FDR < 0.05 were considered. To reduce redundancy among GO terms, one representative term was selected from each hierarchical group displayed in PANTHER, choosing the most specific child term with at least five associated DEGs. Subsequently, terms that were highly redundant were excluded. The resulting GO terms were ranked by fold enrichment, and the top 20 terms were plotted for each line.

### sRNA-seq

For each of the five lines at generation 9, 5th true leaves were collected at 21 DAG from six individual plants and pooled together. The leaf tissue was flash-frozen in liquid nitrogen and stored at −80 °C until use. Total RNA (without depletion of RNAs) was isolated using the mirVana miRNA Isolation Kit. Briefly, frozen tissue was ground to a powder using glass beads using Qiagen TissueLyser II and resuspended in 300 μl of Lysis/Binding Buffer, followed by the addition of 30 μl miRNA Homogenate Additive. RNA was purified with acid-phenol:chloroform and washed in a filter cartridge with 100% ethanol, followed by the mirVana Wash Solution 1 and Wash Solution 2/3. RNA was then eluted with the mirVana Elution Solution, pre-heated to 95°C. 1 μl of Qiagen spike-ins were added to each sample. Libraries were prepared by Novogene and sequenced with NovaSeq 6000. For mapping, sRNA reads were first aligned to Qiagen spike-ins using Bowtie2 [45], and reads that did not align to the spike-ins were retained for genomic mapping. The retained reads were then aligned to the TAIR10 genome using ShortStack v4.1.2 [46]. Known Arabidopsis mature miRNA sequences from miRBase were provided to ShortStack to guide miRNA locus identification. Sequencing and mapping statistics for sRNA-seq libraries are provided in S2 Table.

### Pol V transcript analysis

For Pol V transcription analysis in Figure 4, previously published genomic annotations reflecting Pol V and RdDM activity were used [21]. The whole genome was previously classified into four states using a Hidden Markov Model (HMM), based on the likelihood of Pol V transcription and RdDM activity in each bin [21]. Here, two of the HMM states were combined, and hyperMRs of different classes were overlapped with the resulting classified clusters to obtain Figure 4A. The random control regions were randomly sampled from a set of 300 bp bins which were lowly methylated in generation 2 *drdd* (average mCG < 50%, mCHG < 20%, and mCHH < 5%), with five independent random samplings per context. These controls were selected because they had potential to be classified as a de novo hyperMR by our DMR classifying script. The numbers of control regions were 500, 200 and 200 for CG, CHG and CHH, respectively, approximating average numbers of de novo hyperMRs called in each sequence context.

For read alignments in Figure 4B, IPARE-seq reads from Col-0 samples were obtained from GEO accession GSE146913 [21]. For each read pair, the cDNA-derived sequence was contained in the R1 read, while the unique molecular identifier (UMI) was extracted from the corresponding R2 read using UMI-tools [47] and appended to the matching R1 read name. Poly(A) sequences were trimmed using cutadapt [39], and reads shorter than 20 bp after trimming were discarded. Reads were then aligned as single-end reads to the TAIR10 genome using Bowtie2 [45], allowing one mismatch per seed alignment. Reads with mapping quality ≥ 2 were retained using SAMtools [48], and PCR duplicates were removed using UMI-tools deduplication. For genome browser visualization, deduplicated BAM files from all six Col-0 replicates were merged and indexed using SAMtools [48].

## Acknowledgements

We are most grateful to Andrzej Wierzbicki and Shriya Sethuraman for openly sharing their data and analysis on RNA polymerase V transcripts. We are grateful to Dawei Dai for assistance with EM-seq library preparation and to Ken Chen, Dawei Dai, and Nicole K. Smoot for assistance with bioinformatics analysis.

## AI use disclosure

AI was used in the final proofreading of the manuscript, as well as in the construction of analysis pipelines. The accuracy of all AI output was rigorously checked and verified by the authors.

## Supporting information

S1 Table. Genotyping primers for *drdd*.

S2 Table. Sequencing and mapping statistics for all libraries.

S3 Table. Differentially expressed gene numbers for all comparisons.

S1 Data. *drdd* hyperMRs.

S2 Data. WT hyperMRs.

S3 Data. *drdd* hypoMRs.

S4 Data. WT hypoMRs.

S5 Data. Raw RNA-seq gene counts.

S6 Data. Rlog-transformed RNA-seq gene counts.

S7 Data. DESeq2 differential expression results for *drdd* A vs. 18 WT samples.

S8 Data. DESeq2 differential expression results for *drdd* B vs. 18 WT samples.

S9 Data. DESeq2 differential expression results for *drdd* A vs. 6 WT samples.

S10 Data. DESeq2 differential expression results for *drdd* B vs. 6 WT samples.

---

**S1.**
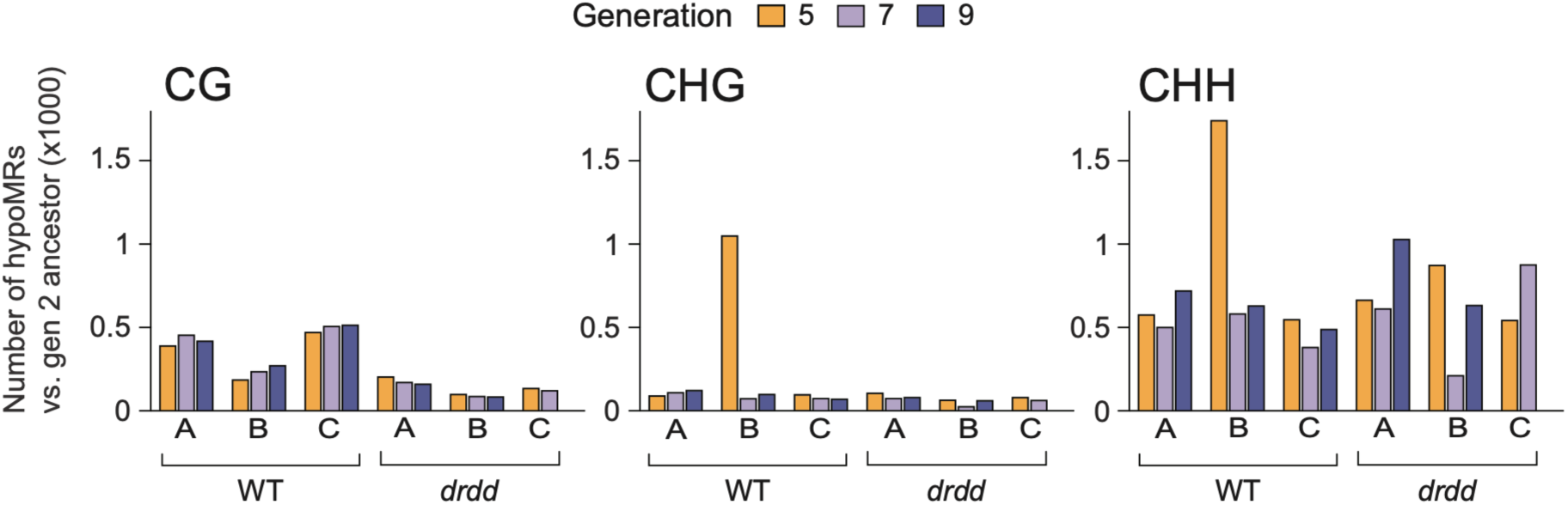
The numbers of hypoMRs in CG, CHG, and CHH sequence contexts in each line relative to their respective generation 2 ancestor, across all analyzed generations.

**S2.**
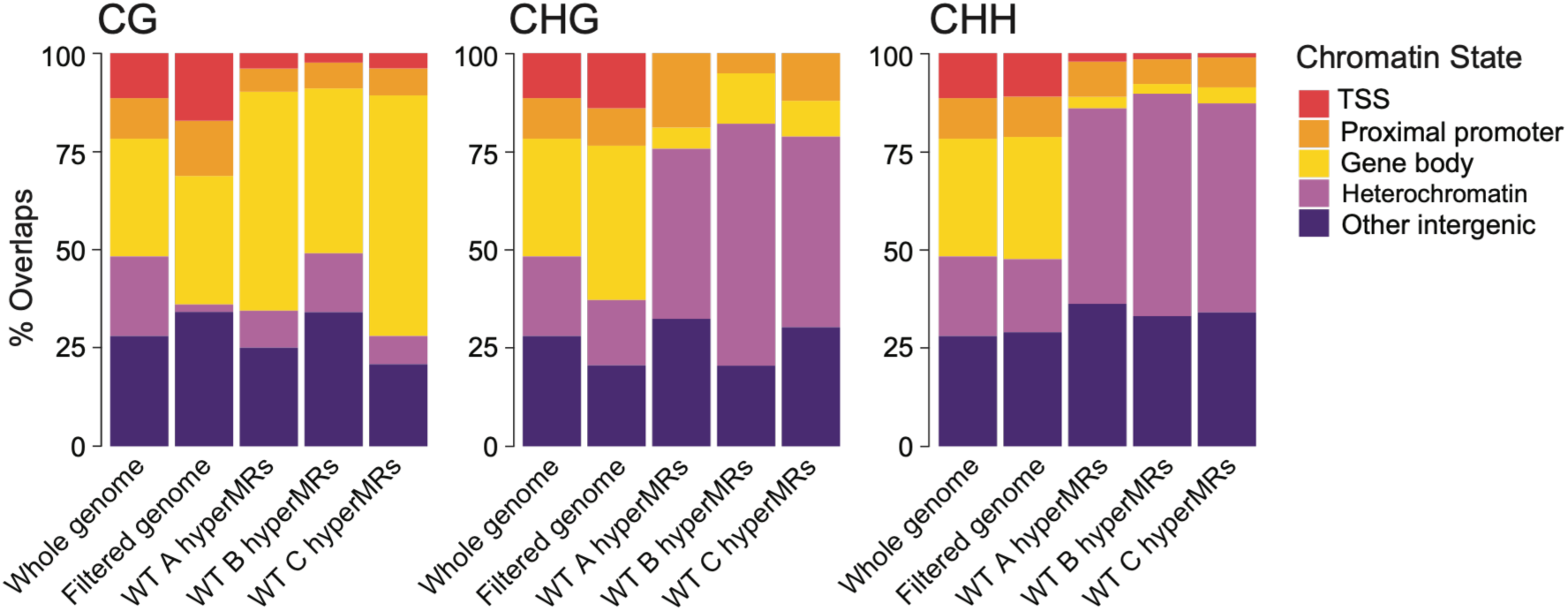
Proportions of WT hyperMRs in generation 7 overlapping major chromatin states (see Methods). “Filtered genome” consists of 300 bp bins which have average methylation below 65% in the CG context, 75% in the CHG context, and 90% in the CHH context in generation 2 WT.

**S3.**
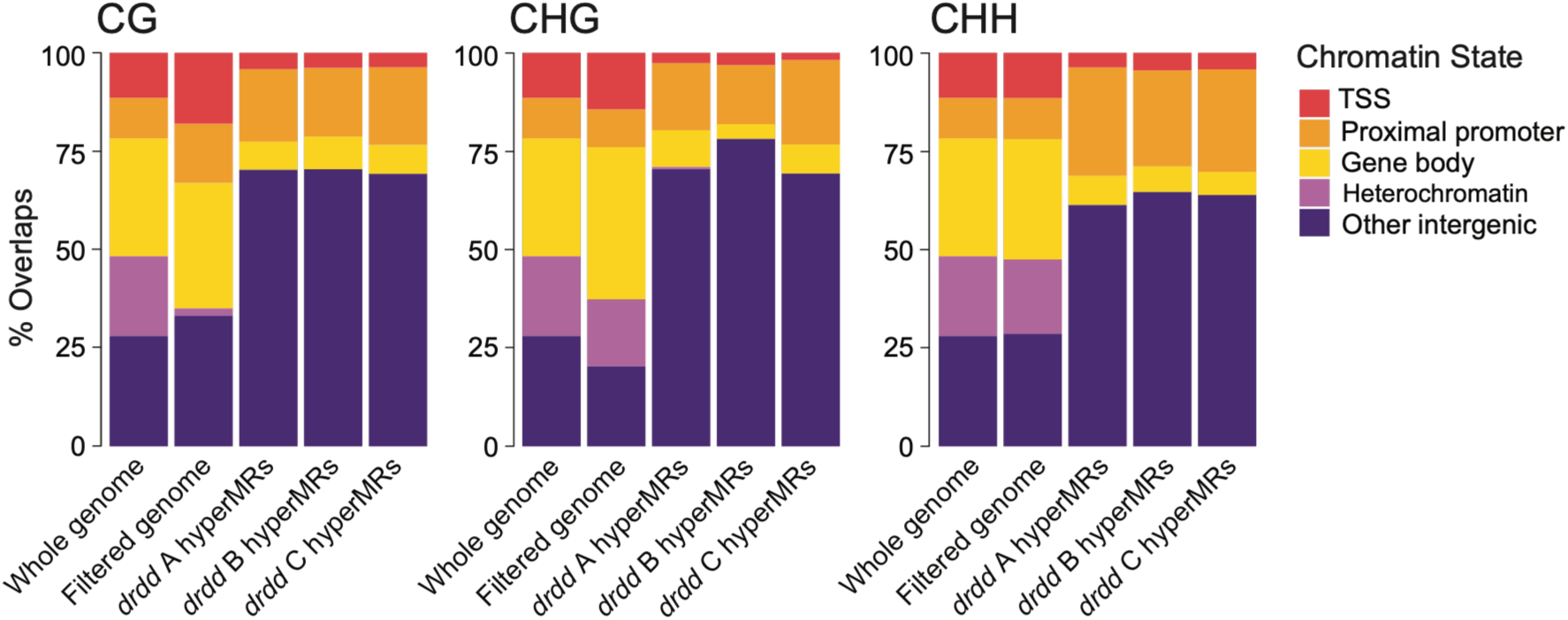
Proportions of *drdd* de novo hyperMRs in generation 7 overlapping major chromatin states. “Filtered genome” consists of 300 bp bins which have average methylation below 65% in the CG context, 75% in the CHG context, and 90% in the CHH context in generation 2 *drdd*.

**S4.**
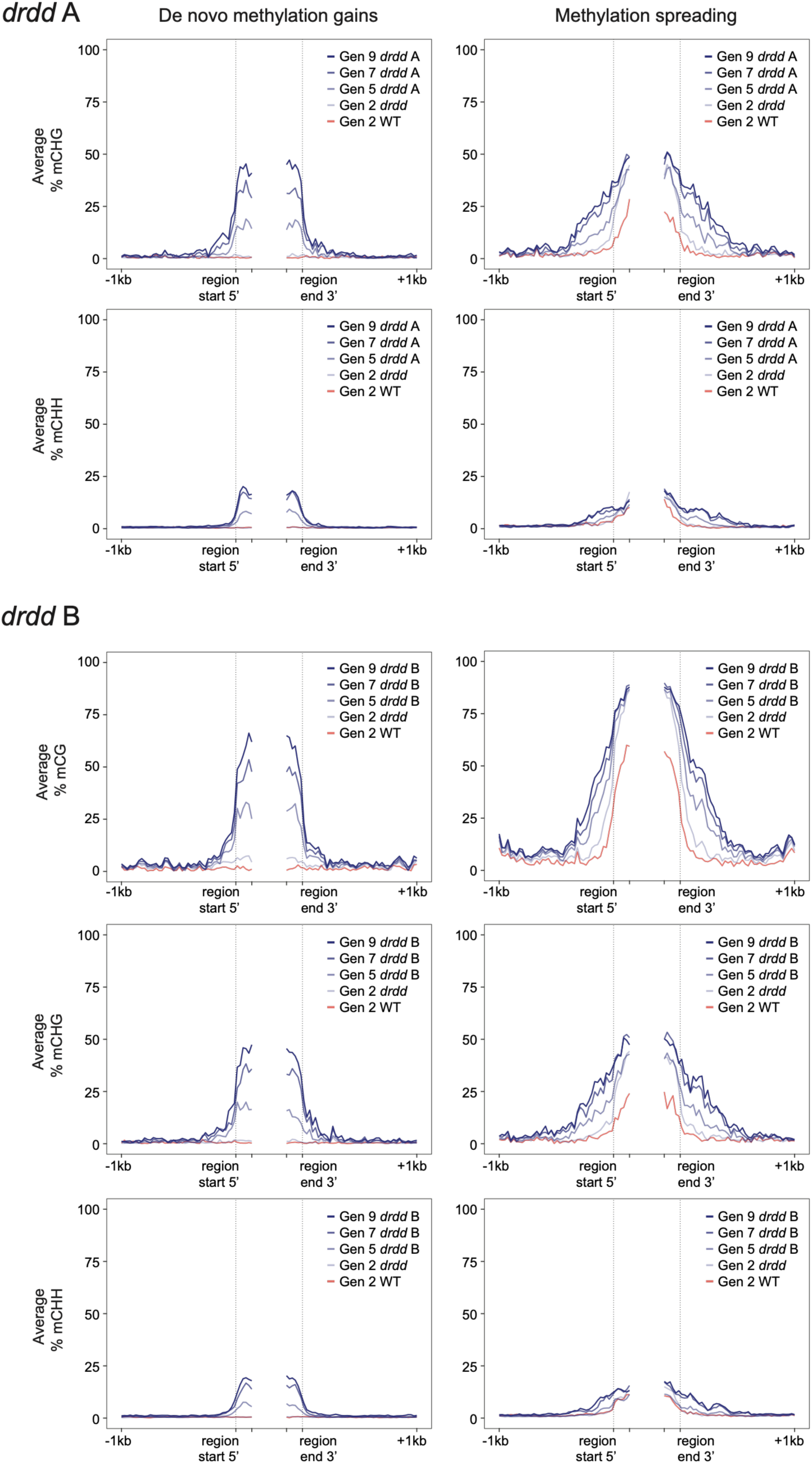
Ends analysis plots showing average methylation levels over de novo and spreading hyperMRs called in generation 9, relative to generation 2 *drdd.* Plots are shown for CHG and CHH contexts in *drdd* line A and all contexts in *drdd* line B. Generation 2 WT is shown for comparison. 150 bp are shown on either side inside the dashed lines, and methylation is averaged over 25 bp bins. Left: de novo hyperMRs are between the vertical dashed lines. Right: spreading hyperMRs are outside of the vertical dashed lines, and the ancestral methylated regions adjacent to them are between the dashed lines.

**S5.**
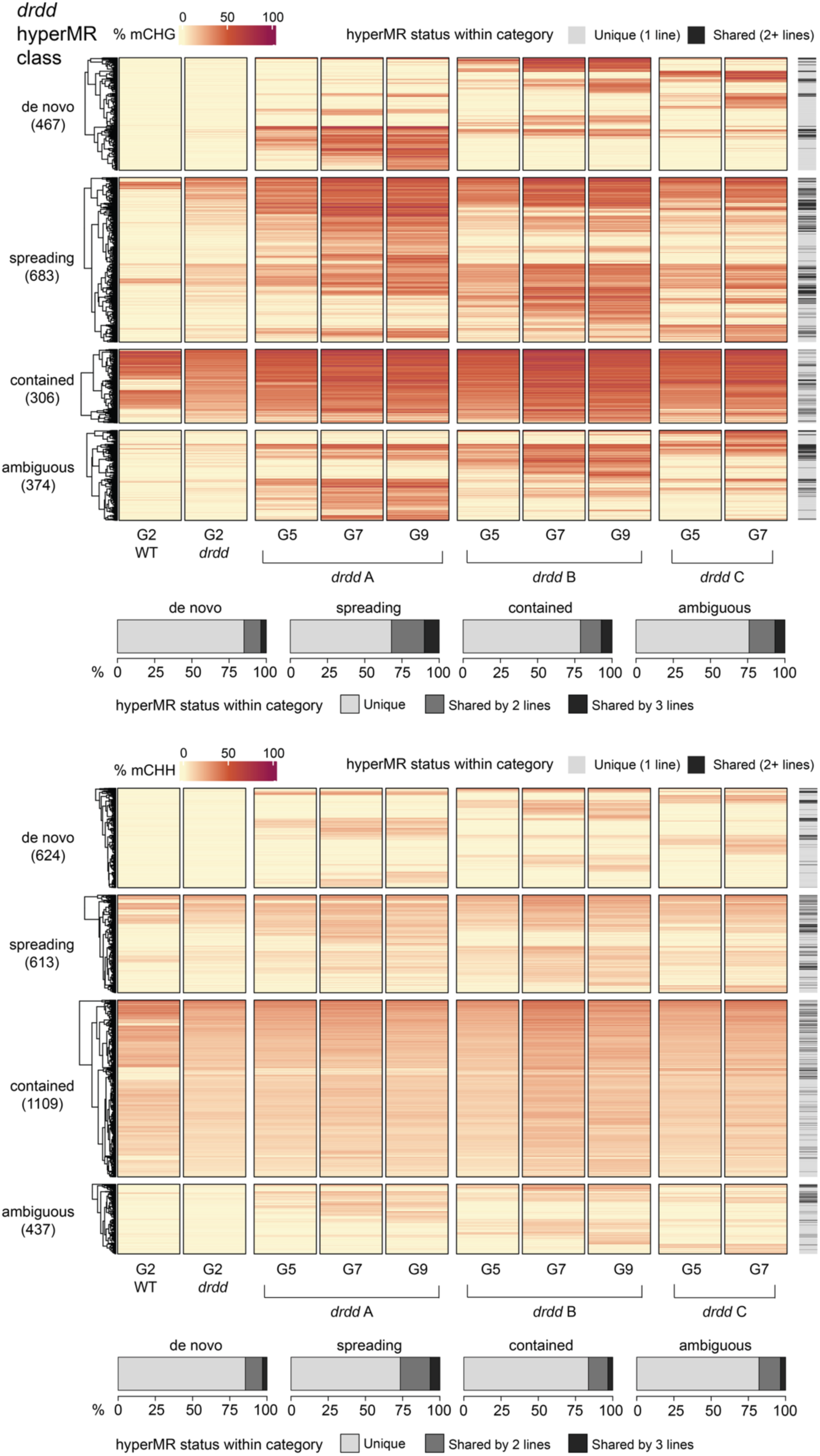
Heatmaps of average methylation in *drdd* lines over class-specific supersets of all *drdd* CHG (above) and CHH (below) hyperMRs ever called, between all lines and generations. Generation 2 WT is shown for comparison. HyperMRs are sorted by hierarchical clustering within each class. Total numbers of hyperMRs in each reduced superset are included in parentheses underneath each hyperMR class name. In the hyperMR status within each class, “Unique (1 line)” means a hyperMR was only called in one line throughout the experiment. “Shared (2+ lines)” means a hyperMR ever called in one line was called in at least one other line in at least one of the generations. Underneath the heatmaps are distributions of *drdd* hyperMRs present in 1 line or shared between 2 or 3 lines in each hyperMR class.

**S6.**
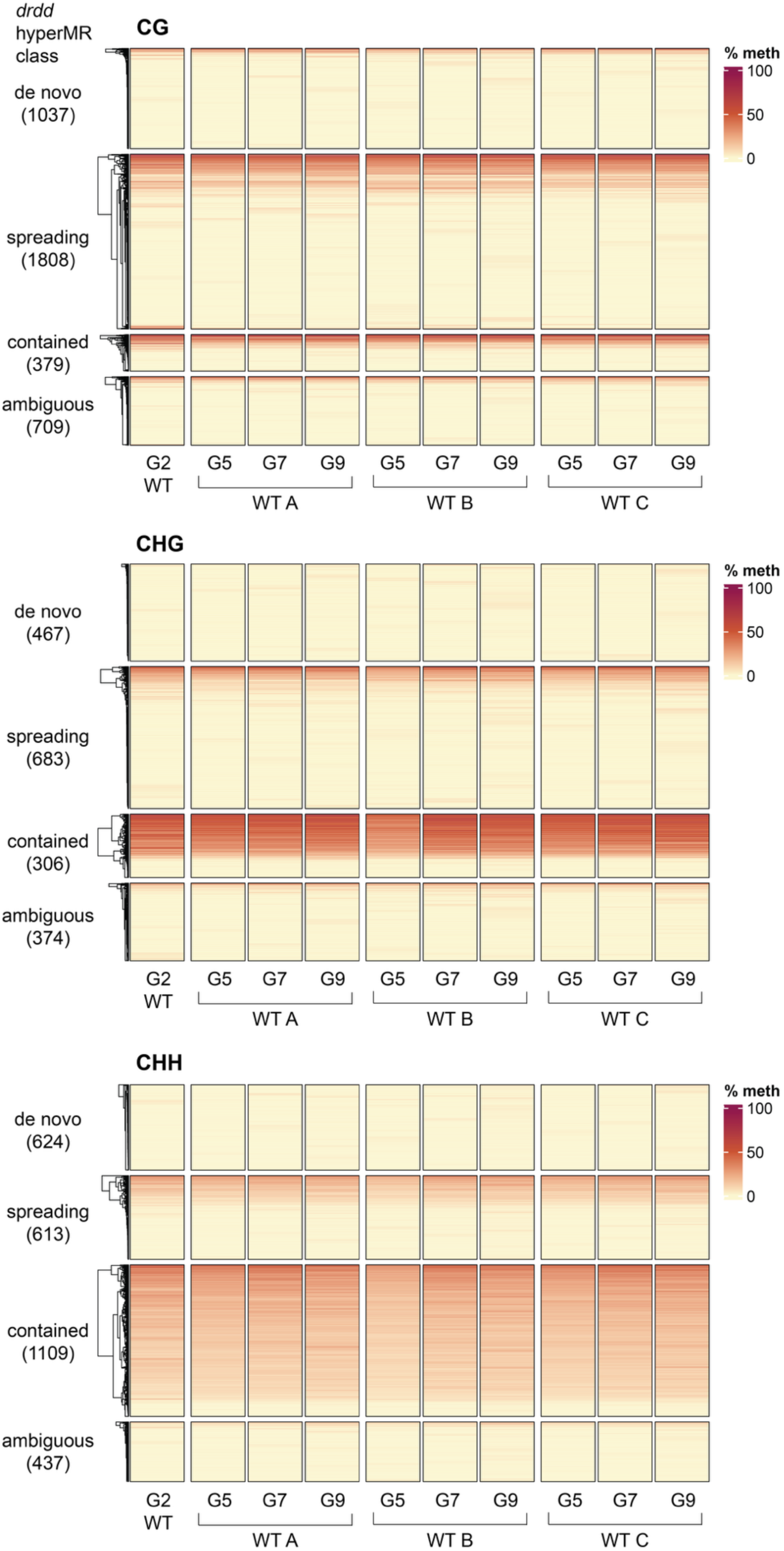
Heatmaps of average methylation in WT lines over class-specific supersets of all *drdd* hyperMRs ever called, between all lines and generations. Heatmaps for all contexts are shown. HyperMRs are sorted by hierarchical clustering within each class. Total numbers of hyperMRs in each reduced superset are included in parentheses underneath each hyperMR class name.

**S7.**
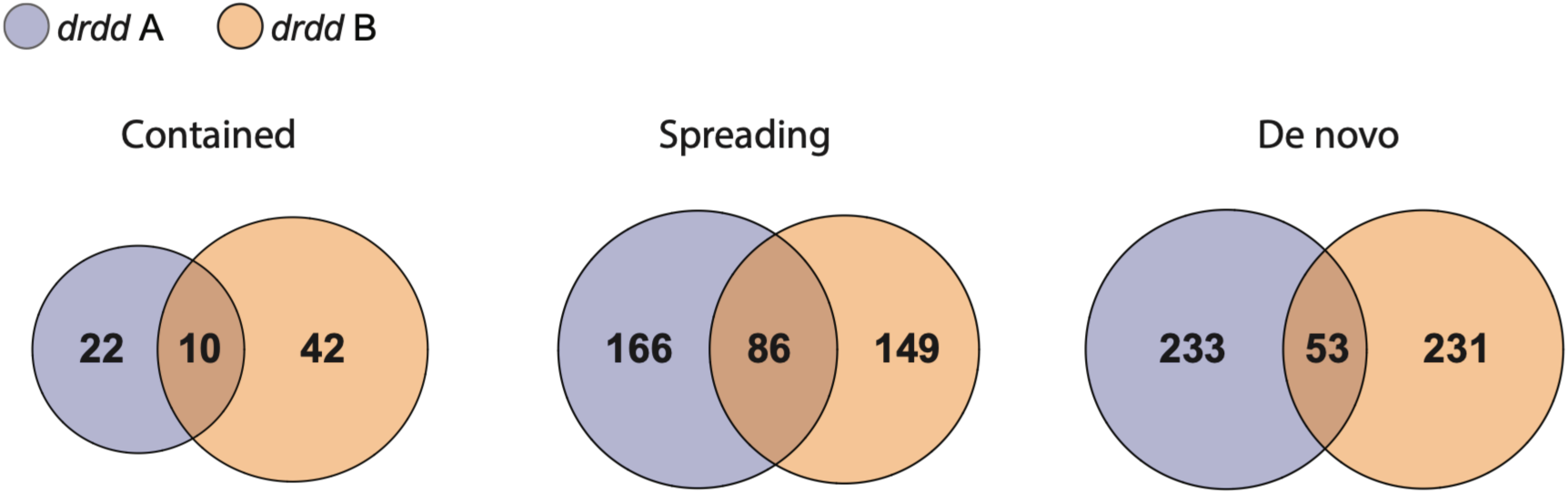
Venn diagrams showing overlap of *drdd-*specific sRNA clusters underlying *drdd* hyperMRs between the two generation 9 *drdd* lines. *drdd*-specific clusters are those that are absent from all WT lines.

**S8.**
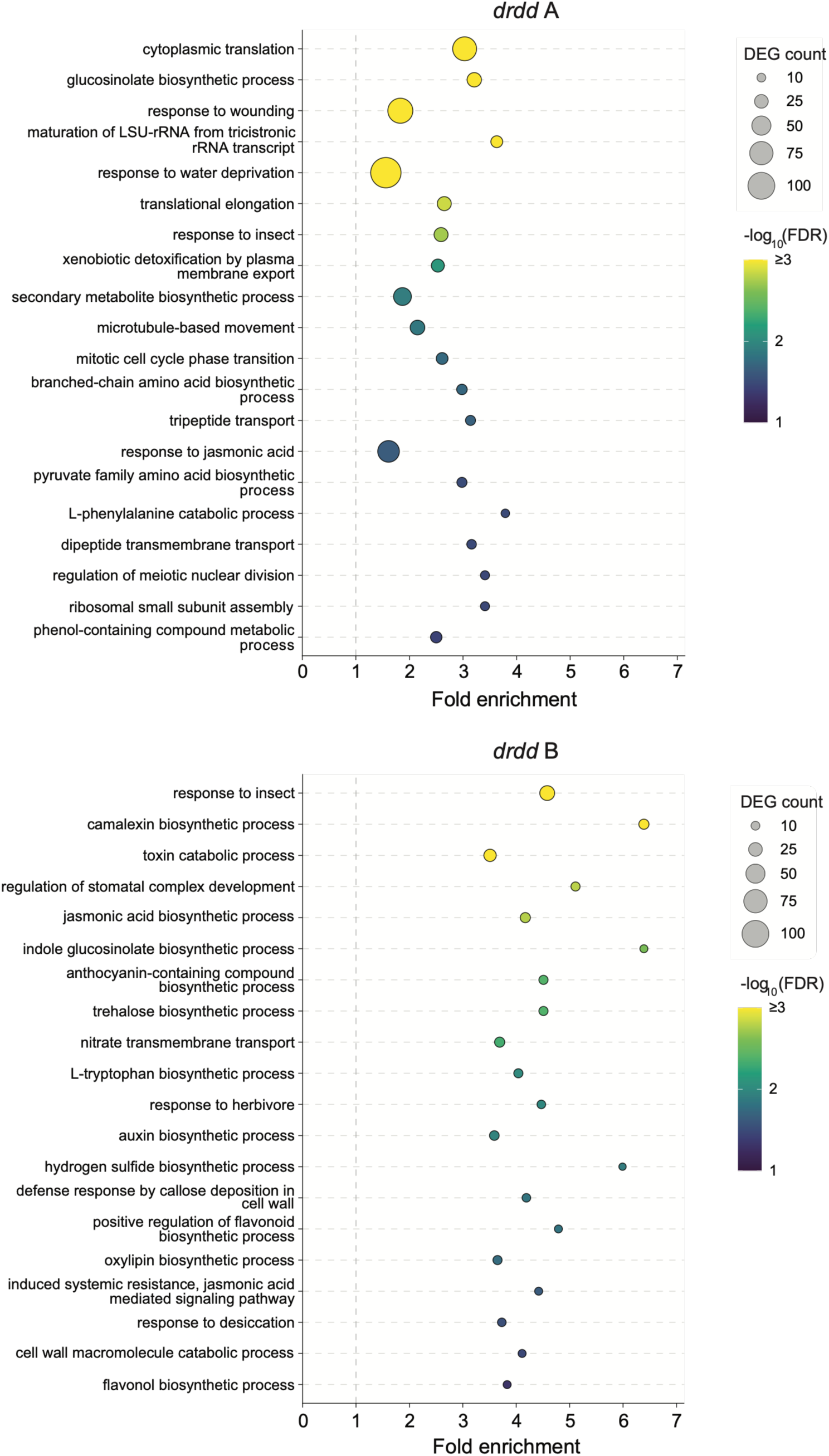
Gene Ontology (GO) enrichment analysis of differentially expressed genes (DEGs) in generation 9 *drdd* lines A and B. For each line, the top 20 GO terms ranked by fold enrichment were selected after reducing redundancy among terms and requiring a minimum of five associated DEGs (see Methods). Bubble size represents the number of DEGs associated with each term, and color indicates −log10(FDR). Some GO term names were slightly abbreviated for clarity where necessary.

